# A novel culture-independent method to search unknown dominant bacterial groups and its application to microbial analysis of membrane bioreactors

**DOI:** 10.1101/2020.04.07.031104

**Authors:** Katsuji Watanabe, Naoto Horinishi

## Abstract

As we could not get numerical information for unknown unculturable microorganisms through conventional culture-independent analysis methods such as next-generation sequencing, or real time PCR, we developed an original culture-independent method, and searched the numerically dominant bacteria in three industrial membrane bioreactors for livestock farms.

Although *Actinobacteria* was the numerically dominant phylum (9.3×10^5^MPN/mL) on 6/August/2014 in the MBR of A farm, when a bacteria with the same genotype to *Arthrobacter* sp. (AF197047; 4.3×10^5^MPN/mL), and those similar to *Burkholderi*a sp. (AB299593; 4.3×10^5^MPN/mL) were the numerically dominant, after about 13 months (24/October/2015) a number of the *Arthrobacter* genotype increased to 930×10^5^ MPN (230 times) and become dominant, and those similar to the *Microbacterium* sp. (AM403628) increased to 92×10^5^MPN, while that of the *Burkholderi*a genotype disappeared. In the other MBR of B farm, bacteria having a similar genotype to *Enshifer* sp. (AB195268, CP000738), or *Shinorhizobium* sp. (AF227755, AB195268), or *Mesorhizobium* sp. (BA000012, Mso.tians29), or *Agrobacterium vitis* (D12795) was dominant on 18/August/2015 (24×10^5^ MPN) and 30/August/2015 (15.5×10^5^ MPN). In the other MBR of C farm (9/October/2015), bacteria having a similar genotype to uncultured *Betaproteobacteria* (AY921864) was dominant (430×10^5^MPN), followed by uncultured bacterium (74×10^5^MPN ; AM268745), and *Mycobacteriaceae* (AB298730), or *Propionibacteriaceae* (AB298731) (7.4 ×10^5^MPN). There was no common bacterial groups among tested three MBRs. Present results indicated that different kinds of homogeneous bacteria were numerically dominant in the three tested membrane bioreactors, where their numbers and ratios were varied with the duration of the driving periods.

**IMPORTANCE:** Although the conventional molecular-based culture independent methods have been used in place of traditional culture-based methods for microbiological research and expanded information of unculturable low-abundance bacterial groups, not all of them were always highly active in the environment and it was difficult to search for microorganisms among them which were highly active and play an important role in the environment. As numerical data of each bacteria might become an important index to know their activity in environment, we had created a novel culture-independent enumeration method for numerically dominant unidentified bacteria. Through the method, we found that different kinds of homogeneous bacteria were numerically dominant in the three tested membrane bioreactors, whose numbers were high enough to affect the performance of the reactor as a single strain. The method was found useful to specify and trace unknown numerically dominant bacterial groups in a culture independent manner.

## INTRODUCTION

Until now, the unculturability of microorganisms was reported to be caused from the following different factors: artificial ones caused by unknown suitable growth condition such as cases in filamentous bacteria in activated sludge (1, 2), or *Dehalococcoides* sp. in soil (3), intrinsic ones caused from low-abundance, and slow growing environmental microorganisms, such as rare biosphere (4, 5), and acquired ones caused from a dormancy state of culturable bacteria (6-10).

As most environmental bacteria were unculturable in growth media, over the past two decades, molecular-based culture independent methods have been used in place of traditional culture-based methods (11, 12), and has expanded information of unculturable bacterial groups in environments (13). In particular, next-generation sequencing (NSG) was expected to become a powerful tool not only to provide genetic information of all the low-abundance microorganisms but also to relate them to their various functions (14).

Among environmental samples, biological wastewater treatment reactors have been widely been studied by using them, based on the social demand that the reactor was an essential facility to purify water by removing excess organics, nitrogen, phosphorus and pathogenic microorganisms in wastewater (15), and that their microbial complex community primarily affected the performance of reactors (16, 17).

As for conventional activated sludge (CAS) equipped with final settling tanks for solid-liquid separation, the culture independent methods such as clone library sequencing (18), denaturing gradient gel electrophoresis (DGGE) (19, 20), terminal restriction fragment polymorphisms analysis (21-23), fluorescence in situ hybridization (FISH) (24, 25), microarray-based genomics (26), and NGS (27-32) have been used to clarify bacterial relation to their performance. These methods have also been used for membrane bioreactor (MBR), by which solids and liquids could be completely separated through a membrane, was an increasingly implemented technology because of advantages such as reduced excess sludge, higher effluent quality and operation under higher biomass concentrations over CAS (23, 24, 30).

Most reports have focused mainly on the comparative study of overall microbial composition (23, 26-28, 32), and their changes in the reactor (29, 30). With respect to the well-known specific gene, or bacteria, their numbers were estimated by using real-time PCR (33-35), or by FISH (24, 25). There have been few reports to show whether numerically dominant specific microorganisms, which might affect the performance of the reactor, existed in the reactor or not until now (1, 2). Higher biomass in effluent, and higher performance of the reactors (16, 20, 22-24, 27, 28, 30, 31) implied that the unculturability of microorganisms in the reactor might mainly be cased from unknown growth condition, and not by low-abundance and slow growing microorganisms, nor by dormant cell.

As microbial numbers might become a simple and useful index to show bacterial activity in the environment, the author has developed an original method to specify and trace unknown dominant microorganisms, in which taxonomies of each bacterial groups were elucidated from the multiple enzyme restriction fragment length polymorphism (MERFLP) of 16S rDNA (36, 37), and each number of the bacterial groups were estimated by most probable number (MPN) (38). Although the method had mostly been applied as a culture-based method (39-43), we presented the results of our application as culture-independent analyses of three industrial membrane bioreactors (MBR) situated in the Hokkaido area. The results were compared with those by the culture-based same method, and those by culture-independent PCR-DGGE.

## MATERIALS AND METHODS

### Samples

The three industrial MBR had been constructed by BICOM Co.Ltd. (Osaka, Japan) to purify and clean up wastewater from daily farms. All the samples were collected from equalizing tanks before septic tanks having an immersion-type membrane filtration apparatus. Sludge in MBR of A farm (Yubetsu Town in Monbetsu-Gun Hokkaido) was sampled on 6/August/2014 and 24/October/2015. That of B farm (Tsurui Village in Akan-Gun Hokkaido) was sampled on 18/ August /2014 and 30/August/2014. That of C farm (Onbetsu Town in Kushiro Branch Office, Hokkaido) was sampled on 9/October/2015. Except for the MBR of A farm on 6/August/2014, when a large amount of waste milk flowed into the MBR because of epidemic bovine mastitis and the performance of the MBR became lower, the total performance of purifying and cleaning up wastewater was kept in good condition.

### Culture-independent MPN and culture based MPN

For culture-independent MPN, DNA was extracted from samples (10mL fresh wt.) as described previously (36) after mixing in a 15 mL Falcon tube (215rpm, 20min). After purification by conventional methods, the DNA solution was further purified using a GenElute Agarose Spin Column (SIGMA). Serial 10-fold dilutions (10^−2^ to 10^−9^) were prepared from the DNA solution.

For culture-based MPN, serial 10-fold dilutions (10^−2^ to 10^−9^) prepared from samples (1mL fresh wt.) were inoculated to test vials (three replicates), including lactose broth (Difco, Sparks MD). After three days incubation at 30°C, bacterial DNA in each vial was extracted as described previously and purified by conventional methods (36, 37).

### MERFLP of the amplified 16S rDNA

Using the V2 forward primer (41f), and the V6 reverse primer (1066r) (44), 16S rDNA was amplified, as described previously (36, 37). Their restriction fragment lengths were measured by microchip electrophoresis systems (MCE-202 MultiNA; Shimadzu Co., Ltd. Kyoto, Japan) after digestion of the PCR product (10μl) using each of the following 4 restriction enzymes: *Hae*III or *Hha* I or *Rsa* I or *Alu* I (10 units, Takara Bio Co. Ltd. Shiga, Japan) in buffer solution (10xLow salt buffer, Takara Bio Co. Ltd.) and 5 folds dilution by de-ionized water, as described previously (36-43).

### Reference database used for the phylogenetic estimation

The reference database used for this research included 30,844 post-amplification sequence files for the 41f/1066r primers, which were mainly re-edited from small subunit rRNA files in RDP II release 9_61 (45) under 5 - bases mismatches in the both in primer annealing sites (36, 37), and consisted of 1,379 bacterial genera, including uncultured and unidentified bacteria (40-43).

### Data processing for multi-template DNA and phylogenetic estimation

As each MPN vial included multi-template DNAs originating from heterogeneous bacteria, the measured MERFL digested from the homogeneous 16S rDNA was selected among the mixed MERFLs digested from the heterogeneous 16S rDNA, as described previously (36, 37). All the theoretical MERFLs originated from the homogeneous 16S rDNA sequence data. The major restriction fragments (RFs) (represented as H in Table 1-3) were those with the highest relative mole concentration (ratio of fluorescent intensity to fragment size). After subtraction of the major RFs from the mixed heterogeneous RFs, the 2nd major RFs were similarly selected (represented as M in Table 1-3). After subtraction of the second major RFs from the remaining heterogeneous RFs, the 3rd major RFs were similarly selected (represented as L in Table 1-3). The similarity between the measured RFLP (A) and the reference RFLP (B) was calculated as described previously (36-43), based on the pairwise distance (*D*_*AB*_) according to Nei and Li (46). The pairwise distance of the MERFLPs (*D*_*ABME*_) was an average of all the *D*_*ABs*_ for used restriction enzymes. Similarity (%) was (1-*D*_*ABME*_) x 100 (Table 1-3). In the phylogenetic estimation, combinations of the three restriction enzymes were used when the identical theoretical MERFL (100% similarity) was not found using the four restriction enzymes. When the identical theoretical MERFL was not found using three restriction enzymes, combinations of the two restriction enzymes was used. If the identical theoretical MERFL (100% similarity) was not found using the two restriction enzymes, the theoretical MERFL having the highest similarity (over 90%) to the measured MERFL was indicated in most cases (Table 1-3) (38-43).

**Table 1.**
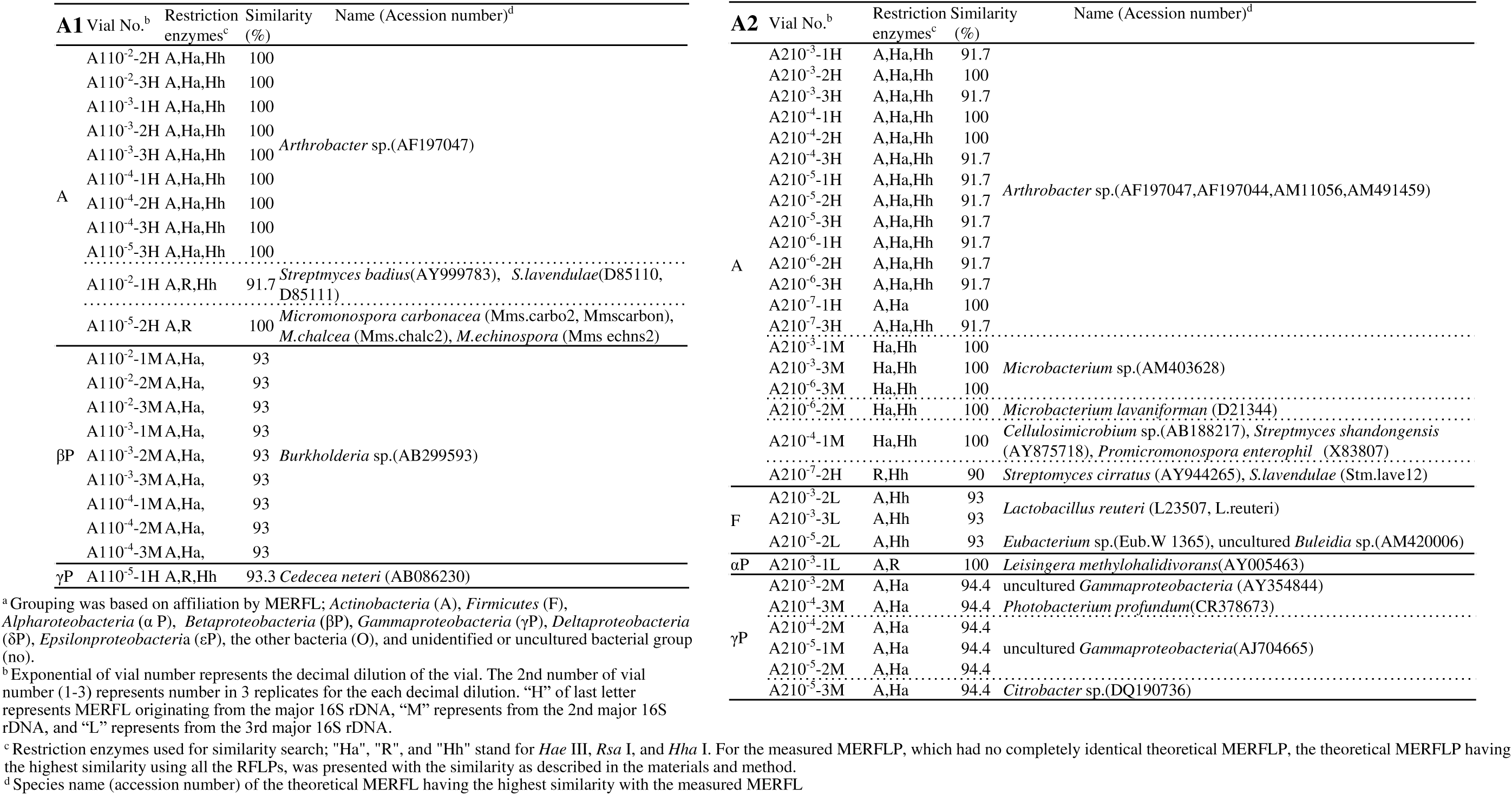
Affiliation^a^ of bacteria in serially diluted DNA extract of MBR in A farm on 6/August/2014 (A1) and on 24/October/2015 (A2) by MERFLP (culture-independent MPN).

**Table 2.**
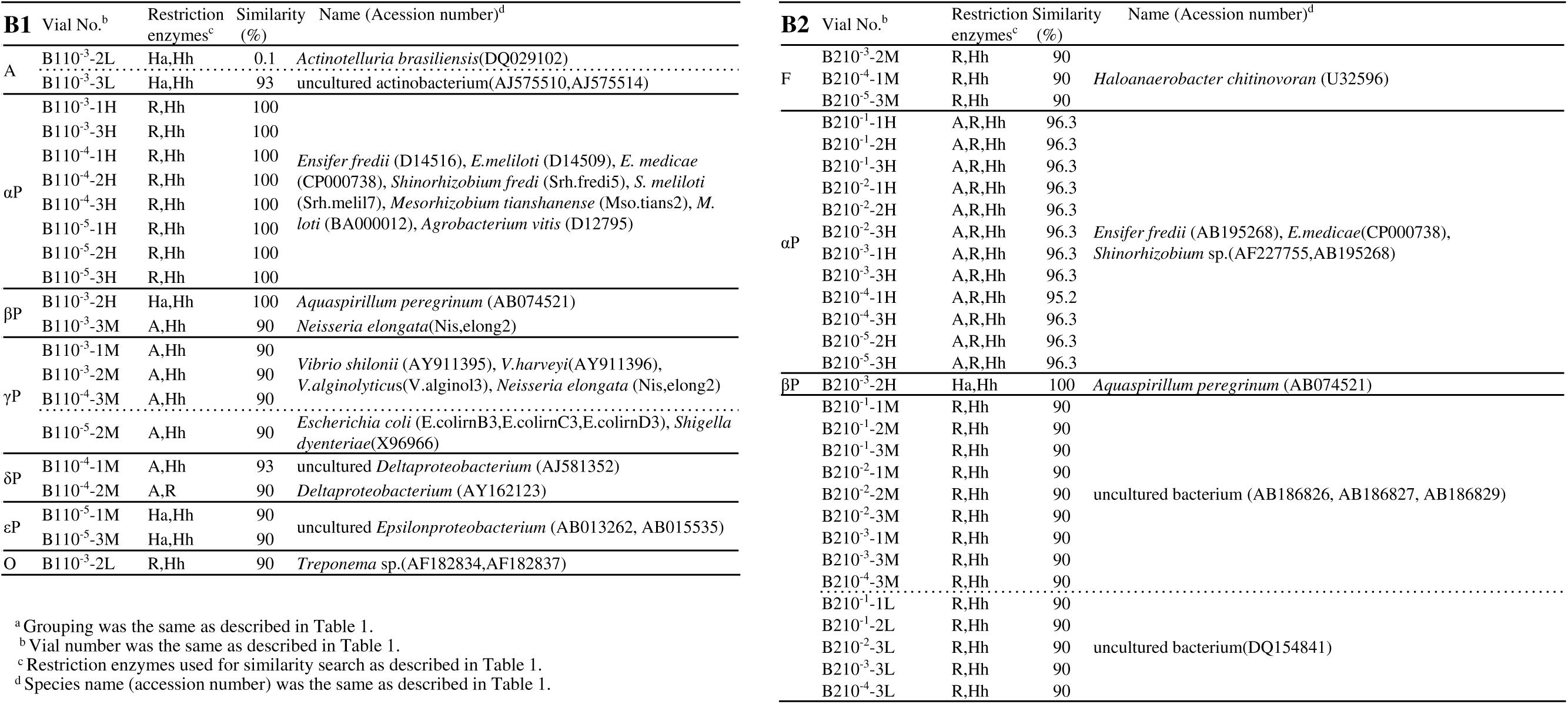
Affiliation^a^ of bacteria in serially diluted DNA extract of MBR in B farm on 18/August/2014 (B2) and on 30/August/2014 (B2) by MERFLP (culture-independent MPN).

**Table 3.**
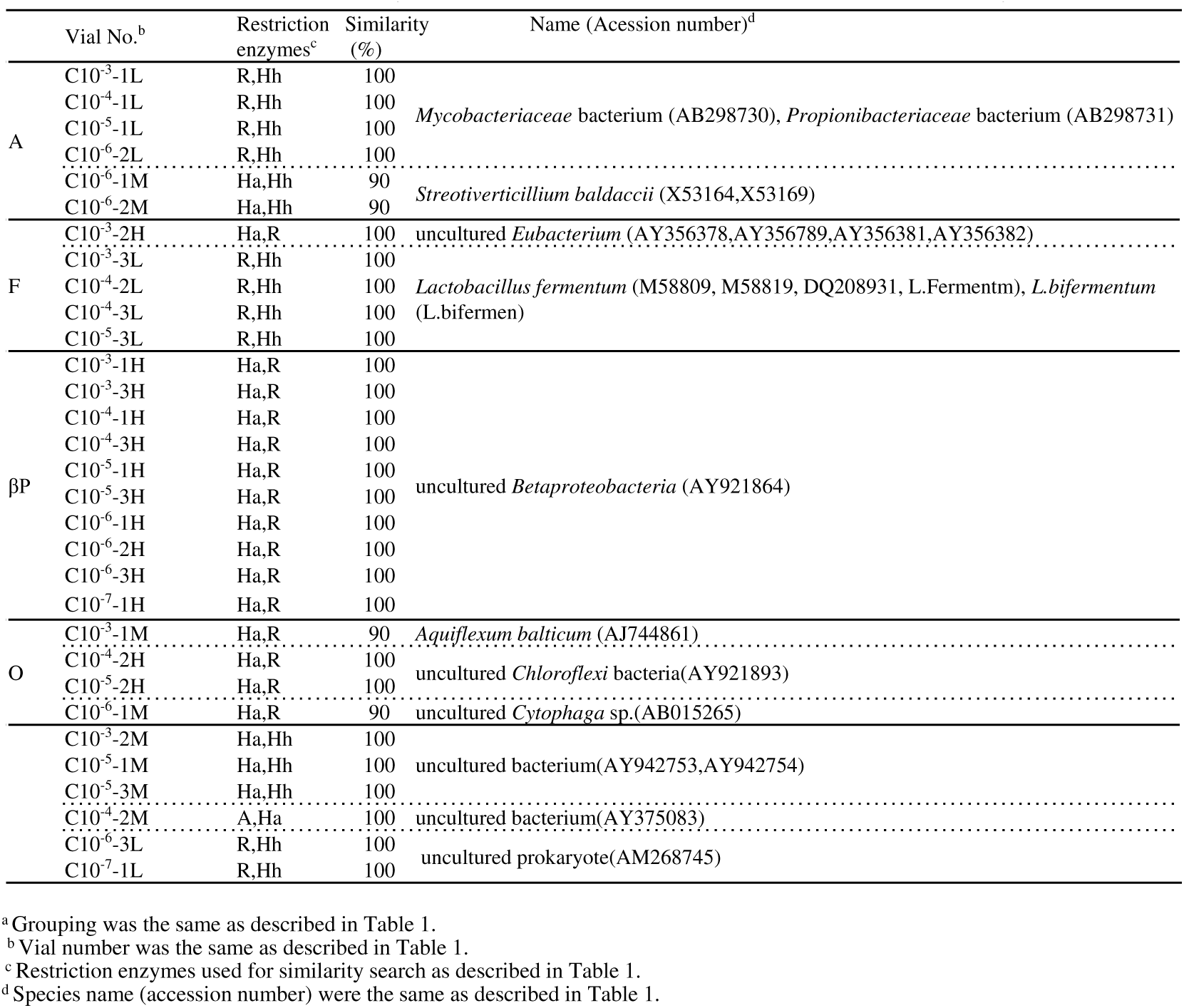
Affiliation^a^ of bacteria in serially diluted DNA extract of MBR in C farm 9/October/2015 by MERFLP (culture-independent MPN).

### Enumeration of bacterial groups by MPN

Through a three-tube, three-decimal-dilution experiment, MPNs of each bacterial groups were estimated (Table 4-6). Using FDA’s Bacterial Analytical Manual (47), confidence limits were obtained and shown in the Tables. Confidence limits shown in Table 4-6 were obtained using Woodward’s method (48), except for the culture-independent MPN in B farm on 30/August/2014, when we could not obtain the data of a 10^−5^ dilution sample, and a 10^−6^ dilution sample.

**Table 4.** Most probable numbers of the numbers of each group in the MBE in A farm on 6/August/2014 by culture-based MPN (a1), on 6/August /2014 (A1) and on 24/October /2015 (A2) by culture-independent MPN and 5% confidence limits obtained using the FDA’s Bacterial Analytical Manual (47, 48).

|  | a1 (culture-based MPN) |  |  |  | A1 (culture-independent MPN) |  |  |  | A2 (culture-independent MPN) |  |  |  |
| --- | --- | --- | --- | --- | --- | --- | --- | --- | --- | --- | --- | --- |
|  | Three dilutions | Score | x10<sup>6</sup>MPN/mL | 5%limits Low-High | Three dilutions | Score | x10<sup>4</sup>MPN/mL | 5% limits Low-High | Three dilutions | Score | x10<sup>6</sup>MPN/mL | 5% limits Low-High |
| *Actinobacteria* | 10<sup>-7</sup>10<sup>-8</sup>10<sup>-9</sup> | 0-0-1 | 30 | 1.5-96 | 10<sup>-4</sup>10<sup>-5</sup>10<sup>-6</sup> | 3-2-0 | 93 | 18-420 | 10<sup>-6</sup>10<sup>-7</sup>10<sup>-8</sup> | 3-3-0 | 240 | 42-1000 |
| *Arthrobacter* sp. |  |  |  |  | 10<sup>-4</sup>10<sup>-5</sup>10<sup>-6</sup> | 3-1-0 | 43 | 9-180 | 10<sup>-6</sup>10<sup>-7</sup>10<sup>-8</sup> |  | 93 | 18-420 |
| *Microbacterium* sp. |  |  |  |  |  |  |  |  | 10<sup>-6</sup>10<sup>-7</sup>10<sup>-8</sup> |  | 9.2 | 1.4-38 |
| *Firmicutes* | 10<sup>-7</sup>10<sup>-8</sup>10<sup>-9</sup> | 2-0-1 | 140 | 36-420 |  |  |  |  | 10<sup>-3</sup>10<sup>-4</sup>10<sup>-5</sup> | 2-0-1 | 0.01 | 0.004-0.042 |
| *αproteobacteria* | 10<sup>-5</sup>10<sup>-6</sup>10<sup>-7</sup> | 2-0-0 | 0.92 | 0.14-3.8 |  |  |  |  | 10<sup>-3</sup>10<sup>-4</sup>10<sup>-5</sup> | 1-0-0 | 0.004 | 0.00002-0.018 |
| *βproteobacteria* | 10<sup>-7</sup>10<sup>-8</sup>10<sup>-9</sup> | 0-1-0 | 30 | 1.5-90 | 10<sup>-4</sup>10<sup>-5</sup>10<sup>-6</sup> | 3-1-0 | 43 | 9-180 |  |  |  |  |
| *Burkholderia* sp. |  |  |  |  | 10<sup>-4</sup>10<sup>-5</sup>10<sup>-6</sup> | 3-1-0 | 43 | 9-180 |  |  |  |  |
| *γproteobacteria* | 10<sup>-7</sup>10<sup>-8</sup>10<sup>-9</sup> | 1-1-0 | 74 | 13-200 | 10<sup>-4</sup>10<sup>-5</sup>10<sup>-6</sup> | 0-1-0 | 11 | 0.15-11 | 10<sup>-4</sup>10<sup>-5</sup>10<sup>-6</sup> | 2-3-0 | 0.29 | 0.09-0.94 |
| *δproteobacteria* |  |  |  |  |  |  |  |  |  |  |  |  |
| *εproteobacteria* |  |  |  |  |  |  |  |  |  |  |  |  |
| Other | 10<sup>-7</sup>10<sup>-8</sup>10<sup>-9</sup> | 3-1-0 | 430 | 90-1800 |  |  |  |  |  |  |  |  |
| unidentified | 10<sup>-4</sup>10<sup>-5</sup>10<sup>-6</sup> | 1-0-0 | 0.04 | 0.002-0.18 |  |  |  |  |  |  |  |  |
| Total number | 10<sup>-7</sup>10<sup>-8</sup>10<sup>-9</sup> | 3-3-3 | >11,000* | 420- | 10<sup>-4</sup>10<sup>-5</sup>10<sup>-6</sup> | 3-3-0 | 240 | 42-1000 | 10<sup>-6</sup>10<sup>-7</sup>10<sup>-8</sup> | 3-3-0 | 240 | 42-1000 |

**Table 5.**
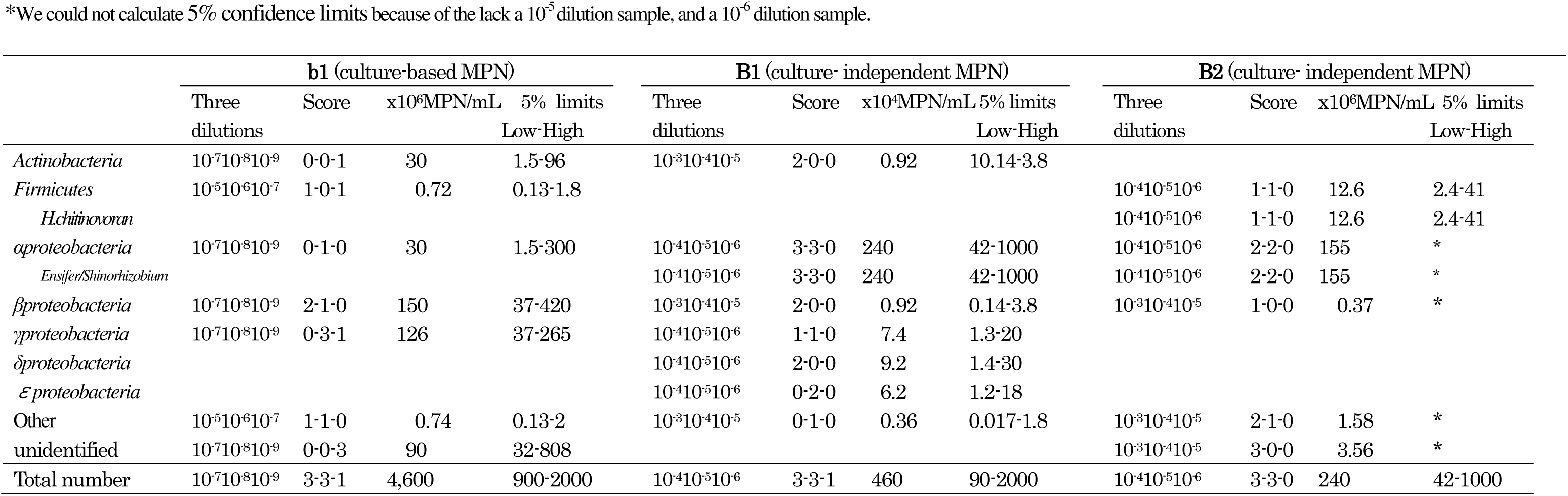
Most probable numbers of the numbers of each group in the MBE in B farm on 18/August/2014 by culture-based MPN (b1), on 18/August/2014 (B1) and on 30/August/2014 (B2) by culture-independent MPN and 5% confidence limits obtained using the FDA’s Bacterial Analytical Manual (47, 48).

**Table 6.** Most probable numbers of the numbers of each group in the MBE in farm C on 9/October/2015 by culture-based MPN (c), and by culture-independent MPN (C) and 5% confidence limits obtained using the FDA’s Bacterial Analytical Manual (47, 48).

|  | c (culture-based MPN) |  |  |  | C (culture-independent MPN) |  |  |  |
| --- | --- | --- | --- | --- | --- | --- | --- | --- |
|  | Three dilutions | Score | x10<sup>6</sup>MPN/mL | 5%limits Low-High | Three dilutions | Score | x10<sup>6</sup>MPN/mL | 5%limits Low-High |
| *Actinobacteria* |  |  |  |  | 10<sup>-5</sup>10<sup>-6</sup>10<sup>-7</sup> | 1-3-0 | 1.6 | 0.45-4.2 |
| Mycobacteriaceae |  |  |  |  | 10<sup>-5</sup>10<sup>-6</sup>10<sup>-7</sup> | 1-1-0 | 0.74 | 0.13-2 |
| *S. baldacii* |  |  |  |  | 10<sup>-5</sup>10<sup>-6</sup>10<sup>-7</sup> | 0-2-0 | 0.62 | 0.12-1.8 |
| *Firmicutes* |  |  |  |  | 10<sup>-4</sup>10<sup>-5</sup>10<sup>-6</sup> | 2-1-0 | 0.15 | 0.037-0.42 |
| *L. fermentum* |  |  |  |  | 10<sup>-4</sup>10<sup>-5</sup>10<sup>-6</sup> | 2-1-0 | 0.15 | 0.037-0.42 |
| *aproteobacteria* |  |  |  |  |  |  |  |  |
| βproteobacteria | 10<sup>-5</sup>10<sup>-6</sup>10<sup>-7</sup> | 0-2-3 | >1.56* | 0.41-2.39 | 10<sup>-6</sup>10<sup>-7</sup>10<sup>-8</sup> | 3-1-0 | 43 | 9-180 |
| AY921864 |  |  |  |  | 10<sup>-6</sup>10<sup>-7</sup>10<sup>-8</sup> | 3-1-0 | 43 | 9-180 |
| γproteobacteria | 10<sup>-5</sup>10<sup>-6</sup>10<sup>-7</sup> | 2-1-0 | 1.5 | 0.37-4.3 |  |  |  |  |
| δproteobacteria |  |  |  |  |  |  |  |  |
| εproteobacteria |  |  |  |  |  |  |  |  |
| Other |  |  |  |  | 10<sup>-5</sup>10<sup>-6</sup>10<sup>-7</sup> | 1-1-0 | 0.74 | 0.13-2 |
| unidentified | 10<sup>-5</sup>10<sup>-6</sup>10<sup>-7</sup> | 1-0-0 | 0.36 | 0.017-1.8 | 10<sup>-6</sup>10<sup>-7</sup>10<sup>-8</sup> | 1-1-0 | 7.4 | 1.3-20 |
| AM26874 |  |  |  |  | 10<sup>-6</sup>10<sup>-7</sup>10<sup>-8</sup> | 1-1-0 | 7.4 | 1.3-20 |
| Total number | 10<sup>-5</sup>10<sup>-6</sup>10<sup>-7</sup> | 3-3-3 | >110* | 42- | 10<sup>-6</sup>10<sup>-7</sup>10<sup>-8</sup> | 1-1-0 | 93 | 18-420 |

### PCR-DGGE

F984GC corresponding to positions 968–984 in *E*.*coli* 16S rRNA (49), and R1378 corresponding to positions 1379-1401 were used as PCR primers (50). The PCR profile consisted of a 2 min denaturation at 94°C, and 30 cycles of 30 sec denaturation at 94°C, a 1 min annealing at 55°C, and a 1 min extension at 72°C, followed by a 3 min extension at 72°C. Amplicons, which were checked by agarose gel electrophoresis, were analyzed by DGGE using a Bio-Rad DCodeTMsystem (Bio-Rad Laboratories, Hercules, CA, USA) according to the manufacture’s manual, as described in the report (51).

## RESULTS

### Culture independent analysis

There was not so large a difference among three replicate electropherograms, which indicated higher reproducibility of the method (Fig. 1). The band strength of the most RFs had gradually decreased in correlation with dilutions, which suggested each of the bacterial numbers could be estimated by MPN (Fig. 1).

**Figure 1.**
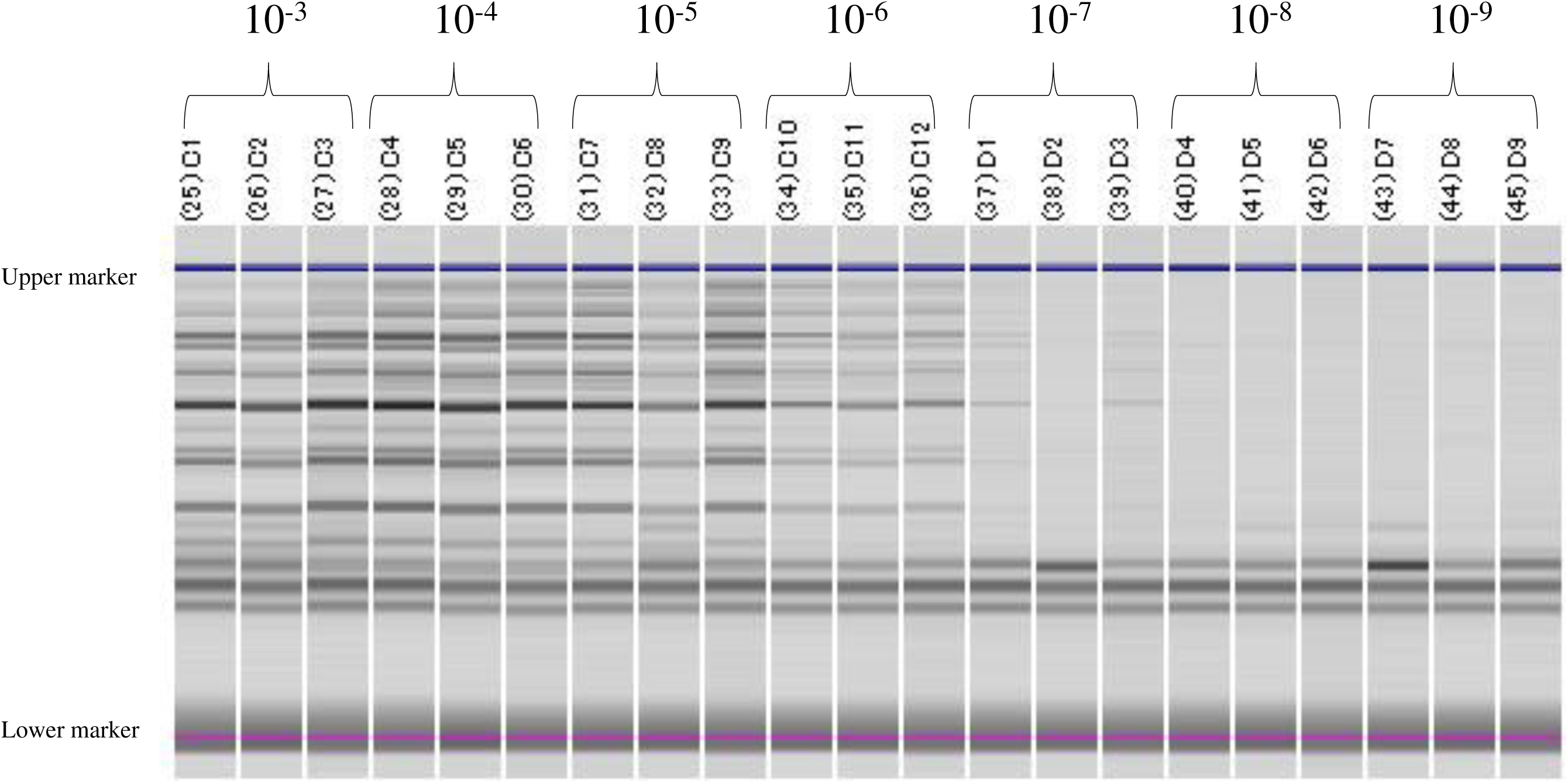
Electropherograms of RFLP (*Hh* I) of serial 10-fold dilutions (10^−3^-10^−9^) in the MBR for C farm on 9/October/2015.

The total detected bacterial number amplified by the 41f/1066r primers was 24×10^5^MPN/mL for MBR in A farm on 6/August/2014. A numerically dominant phylum was *Acitinobacteria* (9.3×10^5^MPN) (Table 1, Table 4), where a homogeneous bacteria with the same MERFLP to *Arthrobacter* sp. (AF197047) was one that was numerically dominant (4.3×10^5^MPN), and that which was similar in MERFLP to *Burkholderia* sp. (AB299593) was the other (4.3×10^5^MPN) (Table 1, 6). After about 13 months (24/October/2015), the total detected bacterial numbers increased to 100 times (2400×10^5^MPN/mL) (Table1, Table 4). All of them were *Actinobacteria*, where those with the *Arthrobacter* sp. genotype were also the numerically dominant (930×10^5^MPN), followed by those with a similar MERFLP to *Microbacterium* sp.(AM403628) (92×10^5^MPN), and those with a *Burkholderia* genotype disappeared (Table 1, Table 4). These results suggested that an increase of the total detected bacteria number during 13 months was caused by a proliferation of *Actinobacteria*, such as the *Arthrobacter* genotype (230 times) and the *Microbacterium* genotype (Fig. 2).

**Figure 2.**
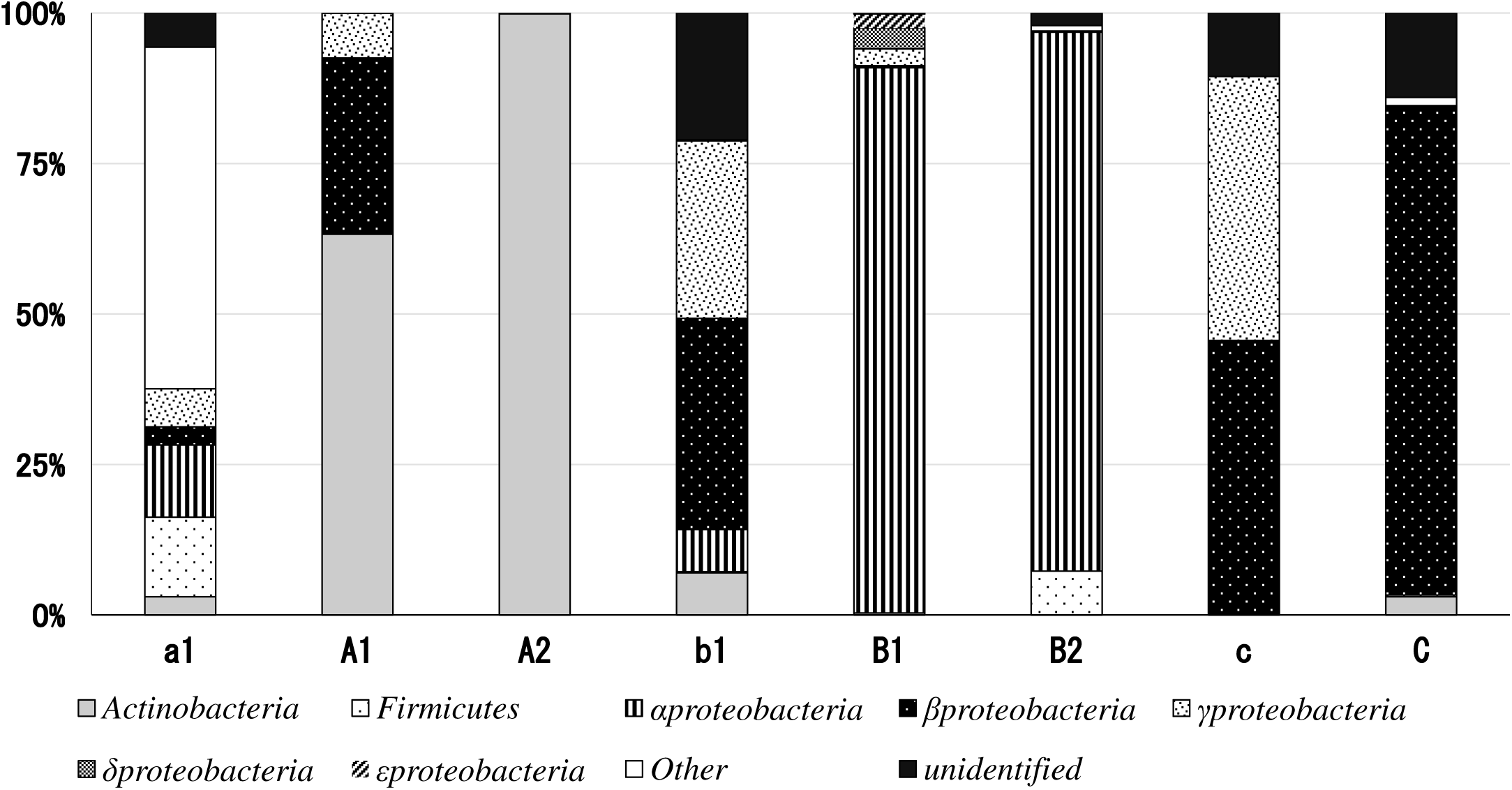
Ratios of bacterial groups estimated by MPN for the tested three MBRs. Capital letters indicate the results of culture-independent MPN as the following: A1, A2 were in the MBR for A farm on 6/August/2014, and 24/October/2015, B1and B2 were the MBR for A farm on 18/ August /2014 and 30/August/2014, and C was the MBR for C farm on 9/October/2015. Small letters indicate the results of culture-based MPN as the following, a1 was in the MBR for A farm on 6/August/2014, b1 was the MBR for B farm on 18/ August /2014, and c was in that for C farm on 9/October/2015.Ratio of Actinobacteria 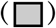, Firmicutes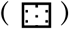, α-Proteobacteria 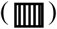, β-Proteobacteria 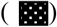, γ-Proteobacteria 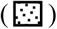, δ-Proteobacteria 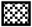),ε-Proteobacteria 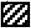), Other bacteria 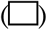, unidentified bacteria 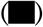.

The total detected bacterial number was 46×10^5^MPN in the other MBR of B farm on 18/August/2015 (Table 5). A numerically dominant phylum was *Proteobacteria*, in which a homogeneous bacteria with a similar MERFLP to *Alphaproteobacteria*, such as *Ensifer* sp.(CP000738,D14509,D14516), or *Shinorhizobium* sp.(Srh.fredi5, Srh.melil7), or *Mesorhizobiu*m sp.(BA000012, Mso.tians2), or *Agrobacterium vitis* (D12795) was numerically dominant (24×10^5^MPN (Table 2, Table 5)). After 12 days (30/August/2015), the total detected bacterial numbers decreased to 24×10^5^MPN/mL (Fig. 2). Although those with the *Alphaproteobacteria* genotype were also numerically dominant, its number decreased to 15.5×10^5^MPN, and a number of bacteria having a similar MERFLP to *Haloanaerobacter chitinovoran* (U32596) increased (4.1×10^5^MPN) (Table 2, Table 5). The result implied that a decrease of the total number of detected bacteria might be caused by a decrease of the numerically dominant *Alphaproteobacteria* genotype (Fig. 2).

The total number of detected bacterial amplified by the 41f/1066r primers was 930×10^5^MPN in the other MBR of C farm on 9/October/2015 (Table 3, 6). A numerically dominant phylum was *Proteobacteria*, in which a homogeneous bacteria having a similar MERFLP to uncultured *Betaproteobacteria* (AY921864) was numerically dominant (430×10^5^ MPN), followed by uncultured bacterium (74×10^5^ MPN; AM268745), *Mycobacteriaceae* (AB298730), or *Propionibacteriaceae* bacterium (7.4×10^5^MPNA; B298731), *Streotiverticillium baldaccii* (6.2×10^5^MPNA;X53164, X53169), and *Lactobacillus fermentum* (1.5×10^5^MPN; DQ208931, L.fermentm) (Table 3, 6). There was no bacteria with the same MERFLP among the three tested MBRs, which implied that the three MBRs had a different bacterial consortium (Fig. 2).

### Comparison to culture-independent PCR-DGGE

There was not such a large difference among three replicate DGGE profiles, which indicated a higher reproducibility of the method (Fig.3, Fig.4). The strength of each bands had gradually decreased in correlation with dilutions, which suggested the numbers of each bands could be estimated by MPN (Fig.3, Fig.4). DGGE profile of MBR in A farm on 6/August/2014 indicated the existence of the two numerically dominant bacteria (B1, and C1), and two subdominant bacterial groups (A1, and D1) (Fig.3). Numbers of B1 and C1 were estimated to be over 11×10^5^MPN, and those of A1 and D1 were 2.3×10^5^MPN (Fig.3). The DGGE profile of MBR in A farm on 24/October/2015 indicated the existence of one numerically dominant bacteria (E2), a subdominant bacteria (D2), and the three bacterial groups (A2, B2, and C2) (Fig.4). The number of E2 was estimated to be over 11,000×10^5^MPN, with that of D2 being 2,300×10^5^MPN. Those of A2, B2, and C2 were 230×10^5^, 230×10^5^, and 23×10^5^, respectively (Fig.4).

**Figure 3.**
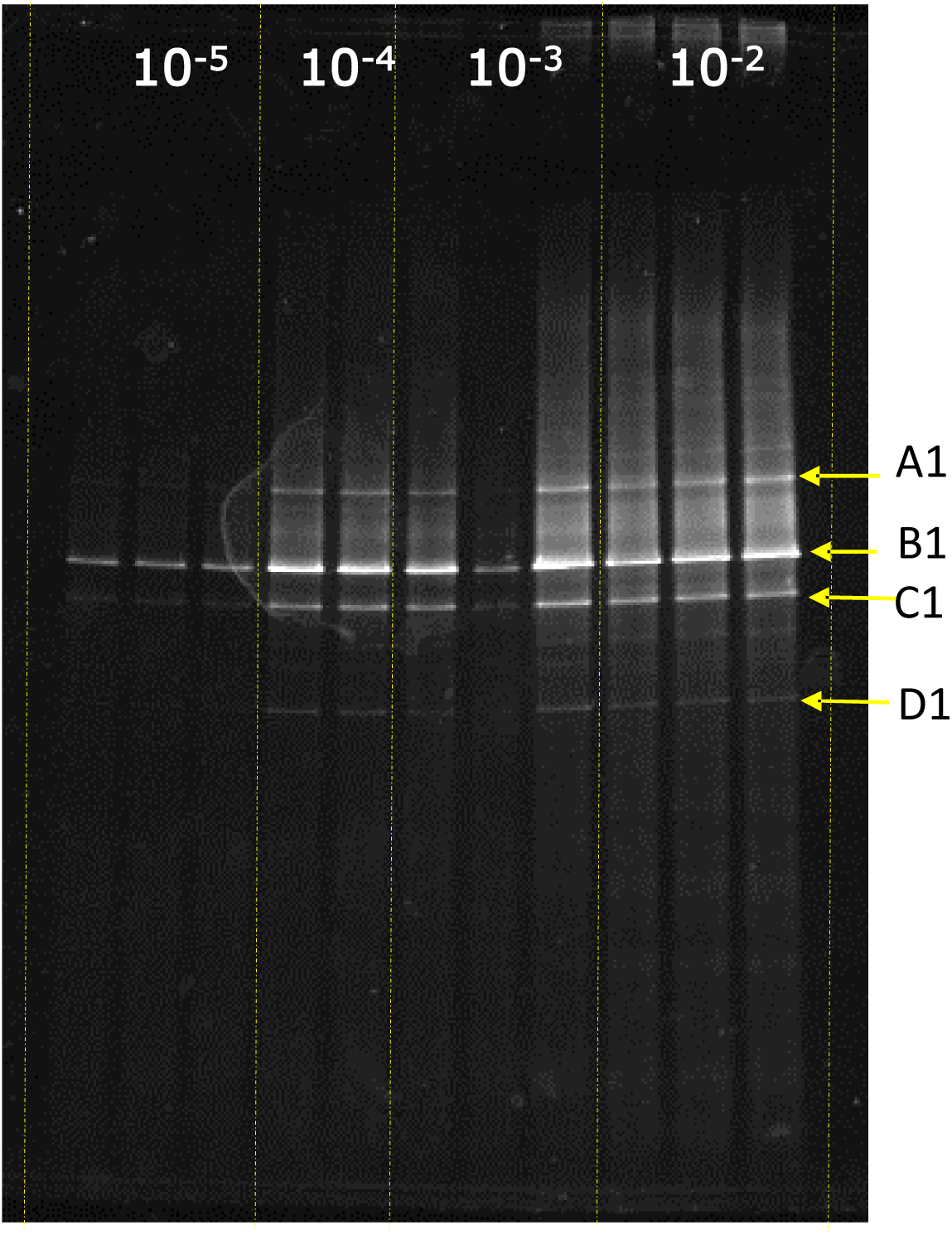
DGGE profiles of serial 10-fold dilutions (10^−2^-10^−5^) in the MBR for A farm on 6/August/2014.

**Figure 4.**
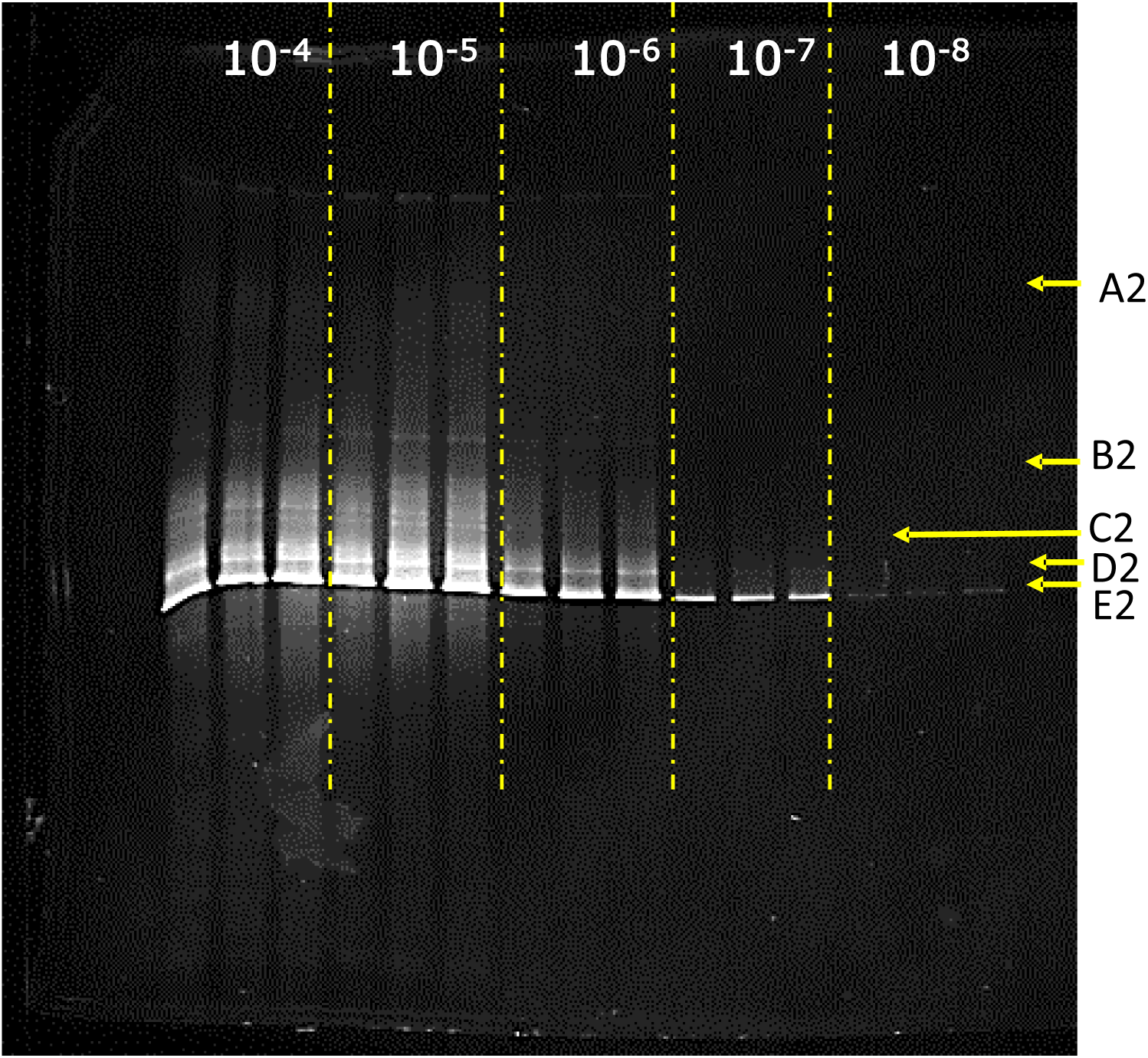
DGGE profiles of serial 10-fold dilutions (10^−4^-10^−8^) in the MBR for A farm on 24/October/2015.

### Comparison to culture-based MPN

As there was a large difference among three replicate electropherograms, this indicated poor reproducibility of the method. Most of the restriction fragments had not always disappeared in correlation with dilutions, which suggested each bacterial number could not precisely be estimated by MPN.

A total detected bacterial number amplified by the 41f/1066r primers was over 110×10^8^MPN/mL for MBR in farm A on 6/August/2014 (Table 4), and that of farm B on 24/October/2015 was 46×10^8^MPN/mL (Table 5). However, the numbers of total bacteria, and the numerical dominant for MBR in farm C on 9/October/2015 were underestimated because the bacterial numbers scaled out of the detection range of the MPN. A total detected bacterial number was over 1.1×10^8^MPN/mL (Table 6). There were no common MERFLPs between those of the culture independent method and the culture-based method (Fig. 2).

## DISCUSSION

All the results of culture-independent MPN indicated that each sample included a homogeneous MERFP (Table 1-Table 3) originated from a single strain (Table 4-Table 6), whose numbers were high enough to affect performance of the reactor (33). However the total number of the detected bacteria was lower than those by the culture-based MPN (Table 4-Table 6), and those by the other reports (16, 33).

The lower bacterial number by the culture-independent MPN was attributed to the elimination of low-abundance bacteria with huge diversity, as the following. There were smeared bands in DGGE profiles lower than 10^−4^ dilution in Figure 3, and lower than 10^−6^ dilution in Figure 4, which were unenumerable diverse kinds of 16S rDNA from low-abundance bacteria, such as rare biosphere (4, 5, 19). They were eliminated from the total bacterial count by our culture independent MPN.

In culture-based MPN, some of such low-abundance bacteria occasionally proliferated in the LB medium, which resulted in the higher total bacterial count (Table 4-6) comparable to those of the reported bacterial numbers (16, 33). However, such an occasional proliferation resulted in incubation bias and poor reproducibility for composition analysis (11, 12), and resulted in a non-detection of the numerically dominant bacteria. LB medium was not suitable for the numerically dominant bacteria in the MBRs. By using growth media suitable for target microorganisms (40, 42), or those which required higher selection bias, such as multi-drug resistant bacteria (38, 43), the culture-based MPN afforded reproducible results.

The results of culture-independent DGGE were well consistent with those of culture-independent MPN. The two numerically dominant bacteria (B1, and C1) in the DGGE profile (Fig.3) were supposed to be the *Arthrobacter* genotype and the *Burkholderia* genotype. The one numerically dominant bacteria (E2) in the DGGE profile (Fig.4) was supposed to be the *Arthrobacter* genotype, and the subdominant bacteria (D2) was supposed to be the *Microbacterium* genotype (Fig.4). Estimated numbers of each identified bacterial groups by our method were lower than those estimated from band patterns in DGGE profiles. Because short RF that originated from higher dilution samples only had lower intensity, it became difficult to select the RFs that originated from homogeneous 16S rDNA precisely, which lowered similarity in the similarity search by MERFLP.

Although the highest number of the *Arthrobacter* genotype was estimated to be 930×10^5^MPN/mL by MERFLP/MPN in the MBR for A farm on 24/October/2015 (Table 4), its number might be underestimated and estimated to be over 11,000×10^5^MPN mL by DGGE/MPN (Fig. 4), which is comparable to the reported total bacteria number in the CAS (from 10,100×10^5^ to 80,000×10^5^cell/mL) (16, 33). The numerically dominant bacteria might primarily influence the performance of the MBR as a single strain.

Although we could not get any information about low-abundance bacteria by our method, the method was found to be effective in specifying and tracing the numerically dominant bacterial groups in the tested three MBRs. This is because precise and reproducible phylogenetic affiliation was possible with respect to the major bacterial groups in higher dilution DNA, where MERFLPs originated from almost a single isolated strain.

As far as CAS reactors, Xia et al. suggested that all the reactors contained a core of bacterial phylum with almost identical compositions, where *Proteobacteria* was the largest phylum and *Firmicutes, Actinobacteria*, and *Bacteroidetes* were the subdominant phylum in five CAS reactors (26). In contrast, Takada et al. reported that there was no core of bacterial phylum or similar phylogenetic structure among 12 different MBRs (23). Our present results, that there was no common numerically dominant bacterial groups in the tested three MBR reactors, was consistent with the latter results (Fig. 2). Our other results, that the composition of overall classes was not changed between sampling periods, was also consistent with those by the other conventional techniques (29, 30) (Fig. 2).

As a new finding through our method, which has not been reported by the other conventional techniques (23, 26-30, 32), the present results indicated that different homogeneous bacteria were numerically dominant in all three MBRs individually, whose numbers were high enough to affect the performance of the reactor as a single strain. The numerically dominant bacteria in A farm were the homogeneous *Arthrobacter* genotype and the homogeneous *Burkholderi*a genotype (AB299593) (Table 1a, 1b), while those in B had the homogeneous *Alphaproteobacteria* genotype (Table 2a, 2b), and those in C had the homogeneous *Betaproteobacteria* genotype (Table 3), which occupied a higher ratio among the detected bacteria (Table 6). This finding could be obtained by the differentiation and elimination of low-abundance bacterial groups having huge diversity shown as smeared bands in the DGGE profiles (Fig. 3, Fig.4).

The present results also indicated that the method was effective to demonstrate the population dynamics of unknown bacterial groups without cultivation, as the following. In the MBR of A farm, the number of the numerically dominant *Arthrobacter* genotype increased 230 times, and a number of the *Microbacterium* genotpe increased to become the subdominant strain, while the other numerically dominant *Burkholderia* genotype disappeared during 13 months (Table 4), when the performance of the MBR recovered to normal condition from serious damage by a large effluent of waste milk. In the MBR of B farm, the number of the numerically dominant *Alphaproteobacteria* genotype (24×10^5^ MPN) slightly decreased to 15.5×10^5^MPN, the number of those of the *Haloanaerobacter chitinovoran* genotype increased after 12 days (Table 5). Our present results clearly demonstrated a dynamic transition of microbial composition in an MBR that particular bacteria proliferated and became extinct.

Until now, most of the microbial research of CAS (18-22, 24-29, 31, 32), and MBR (23, 24, 30) has been focused on the exploration of microbial diversity using the conventional culture-independent molecular techniques with aim to cover almost all the low-abundance microorganisms. These approaches seemed to have targeted such bacterial groups as those that appeared as smeared bands in lower dilutions of DGGE profiles (Fig. 3, Fig.4), which seemed to be not suitable to specify and trace numerically dominant bacterial groups. Our method was simple but effective to clarify the dynamic transition of the numerically dominant unknown microbial group in bioreactors.

## ACKNOWLEDGMENTS

All of the BMR samples were provided by BICOM Co. Ltd. (Osaka, Japan). We thank Mr. S. Senkawa, representative director of BICOM Co. Ltd. for sending us samples. We thank Prof. M.Sakai, and Assistant Prof. M. Ikenaga of the lab of soil science at Kagoshima University for providing us with various advice, information, and suggestions about DGGE techniques. We thank Miss Y. Akahoshi, and Mr. T. Momoshima, undergraduate students, for analysis of the samples. We also thank Mr. Patrick Slusar of Fukuoka Institute of Technology for English proofreading. English.

## REFERENCES

1. Layton AC, Karanth PN, Lajoie CA, Meyers AJ, Grefory IR, Stapleton RD, Taylor DE, Sayler GS. 2000. Quantification of *Hyphomicrobium* Populations in Activated Sludge from an Industrial Wastewater Treatment System as Determined by 16S rRNA Analysis. Appl. Environ. Microbiol. 66: 1167–1174.

2. Speirs L, Nittami T, McIlroy S, Schroeder S, Seviour R J. 2009. Filamentous Bacterium Eikelboom Type 0092 in Activated Sludge Plants in Australia Is a Member of the Phylum *Chloroflexi*_ Appl. Environ. Microbiol. 75: 2446–2452.

3. Adrian L, Szewzyk U, Wecke J, Görisch H. 2000. Bacterial dehalorespiration with chlorinated benzenes. Nature 408: 580–583.

4. Pedrós-Alió C. 2012. The rare bacterial biosphere. Ann. Rev. Mar. Sci. 4: 449–466.

5. Sogin ML, Morrison HG, Huber JA, Welch DM, Huse SM, Neal PR, Arrieta JM, Herndl GJ. 2006. Microbial diversity in the deep sea and the underexplored “rare biosphere”. Proc. Natl. Acad. Sci. U.S.A. 103: 12115–12120.

6. Lleò MM, Pierobon S, Tafi MC, Signoreto C, Canepari P. 2000. mRNA detection by reverse transcription-PCR for monitoring viability over time in an Enterococcus faecalis viable but nonculturable population maintained in a laboratory microcosm. Appl. Environ. Microbiol. 66: 4564–4567.

7. Oliver JD. 2010. Recent findings on the viable but nonculturable state in pathogenic bacteria. FEMS Microbiol. Rev. 34:415–425.

8. Chen DMJ, Zhang X, Li A, Li Y, Wang Y. 2007. Retention of virulence in a viable but nonculturable *Edwardsiella tarda* isolate. Appl. Environ. Microbiol. 73: 1349–1354.

9. Quirós C, Herrero M, Garcia LA, Diaz M. 2009. Quantitative approach to determining the contribution of viable-but nonculturable subpopulations of malolactic fermentation processes. Appl. Environ. Microbiol. 75: 2977–2981.

10. Sachidanandham R, Gin KYH. 2009. A dormancy state in nonspore-forming bacteria. Appl Microbiol Biotec. 81:927–941.

11. Prakasam TBS, Dondero NC. 1967. Aerobic Heterotrophic Bacterial Populations of Sewage and Activated Sludge I. Enumeration’ Appl. Environ. Microbiol. 15:461–467.

12. Dias FF, Bhat JV. 1964. Microbial Ecology of Activated Sludge I. Dominant Bacteria Appl Environ Microb, 12(5):412–417.

13. Rappé MS, Giovannoni SJ. 2003. The Uncultured Microbial Majority. Annu. Rev. Microbiol. 2003. 57:369–94.

14. Fierera N, Leffb JW, Adamsc BJ, Nielsend UN, Batesb ST, Lauberb CL, Owense S, Gilberte J, Wallh DH, Caporasoe JG. 2012. Cross-biome metagenomic analyses of soil microbial communities and their functional attributes. Proc. Natl. Acad. Sci. U.S.A. 109:21390–21395l.

15. Narihiro, T. 2016. Microbes in the water infrastructure: underpinning our society. Microbes Environ. 31:89–92.

16. Hiraishi, A. 1999. Distribution of viable but non-culturable bacteria in wastewater treatment systems. Microbes Environ. 14:91–99. (In Japanease).

17. Kamagata, Y. 2007. What are uncultured Microbe? J. Environ. Biotech. 7:69–73 (In Japanease).

18. Snaidr J, Amann R, Huber I., Ludwig W, Schleifer KH. 1997. Phylogenetic Analysis and In Situ Identification of Bacteria in Activated Sludge. Appl. Environ. Microbiol. 62:2884–2896.

19. Piterina AV, Pembroke JT. 2013. Use of PCR-DGGE Based Molecular Methods to Analyse Microbial Community Diversity and Stability during the Thermophilic Stages of an ATAD Wastewater Sludge Treatment Process as an Aid to Performance Monitoring. ISRN Biotech. 2013: 162645.

20. Hanada A, Kurogi T, Giang NM, Yamada T, Kamimoto Y, Kiso Y, Hiraishi A. 2014. Bacteria of the Candidate Phylum TM7 are Prevalent in Acidophilic Nitrifying Sequencing-Batch Reactors. Microbes Environ. 29: 353–362.

21. Saikaly PE, Stroot PG, Oerther DB. 2005. Use of 16S rRNA Gene Terminal Restriction Fragment Analysis to Assess the Impact of Solids Retention Time on the Bacterial Diversity of Activated Sludge. Appl. Environ. Microbiol. 71:814–5822. t-RFLP AEM 5814 2005

22. V.-Vargas A, T.-Labrador G, M.l-Deya AA 2012. Bacterial Community Dynamics in Full-Scale Activated Sludge Bioreactors: Operational and Ecological Factors Driving Community Assembly and Performance PLoS One 7:e42524.

23. Takeda K, Hashimoto K, Soda S, Ike M, Yamashita K, Hashimoto T. 2014. Characterization of Microbial Community in Membrane Bioreactors Treating Domestic Watewater. J. Water Environ. Tec. 12:99–107.

24. Silva AF, Carvalho G, Oehmen A, L.-Ferreira M, van Nieuwenhuijzen A, Reis MAM., Teresa M, Crespo B. 2012. Microbial population analysis of nutrient removal-related organisms in membrane bioreactors. Appl. Microbiol. Biotechnol. 93:2171–2180.

25. Lenaerts J, Lappin-Scott HM, Porter J. 2007. Improved Fluorescent In Situ Hybridization Method for Detection of Bacteria from Activated Sludge and River Water by Using DNA Molecular Beacons and Flow Cytometry. Appl. Environ. Microbiol. 73:2020–2023.

26. Xia S, Duan L, Song Y, Li J, Andersen GL, Alvarez-Cohen L, Moreno-Andrade I, Huang CL, Hermanowicz S. 2010. Bacterial Community Structure in Geographically Distributed Biological Wastewater Treatment Reactors. Environ. Sci. Technol. 44:7391–7396.

27. Zhang T, Shao MF, Ye L. 2012. 454 Pyrosequencing reveals bacterial diversity of activated sludge from 14 sewage treatment plants. The ISME Journal 6:1137–1147.

28. Wang X., Hu M, Xia Y, Wen X, Ding K. 2012. Pyrosequencing Analysis of Bacterial Diversity in 14 Wastewater Treatment Systems in China. Appl. Environ. Microbiol, 78:7042–7047.

29. Ibarbalz FM, Orellana E, Figuerola ELM, Erijman L. 2016. Shotgun Metagenomic Profiles Have a High Capacity To Discriminate Samples of Activated Sludge According to Wastewater Type. Appl..Environ. Microbiol. 82:5186–5196.

30. Sato Y, Hori T, Navarro RR, Naganawa R, Habe H, Ogata A. 2016. Effects of Organic-Loading-Rate Reduction on Sludge Biomass and Microbial Community in a Deteriorated Pilot-Scale Membrane Bioreactor. Microbes Environ. 31:361–364.

31. Narihiro T, Nobu MK, Hori T, Aoyagi T, Sato Y, Inaba T, Aizawa H, Tamaki H, Habe H. 2019. Effects of the Wastewater Flow Rate on Interactions between the Genus *Nitrosomonas* and Diverse Populations in an Activated Sludge Microbiome. Microbes Environ. 34:89–94.

32. Xia S, Duan L, Song Y, L i J, Piceno YM, Andersen GL, A.-Cohen L, M-Andrade I, Sanapareddy N, Hamp TJ, Gonzalez LC, Hilger HA, Fodor AA, Clinton SM. 2009. Molecular Diversity of a North Carolina Wastewater Treatment Plant as Revealed by Pyrosequencing. Appl. Environ. Microbiol. 75:1688–1696.

33. Dionisi HM, Harms G, Layton AC, Gregory IR, Parker J, Hawkins SA, Robinson KG, Sayler GS. 2003. Power Analysis for Real-Time PCR Quantification of Genes in Activated Sludge and Analysis of the Variability Introduced by DNA Extraction. Appl. Environ. Microbiol. 69:6597–6604.

34. Baptista JDC, Lunn M, Davenport RJ, Swan DL, Read LF, Brown MR, Morais C, Curtisa TP. 2014. Agreement between *amoA* Gene-Specific Quantitative PCR and Fluorescence *In Situ* Hybridization in the Measurement of Ammonia-Oxidizing Bacteria in Activated Sludge. Appl. Environ. Microbiol. 80: 5901–5910.

35. Nguyen VL, He X, de los Reyes III FL. 2016. Quantifying in situ growth rate of a filamentous bacterial species in activated sludge using rRNA:rDNA ratio. FEMS. Microbiol. Letters. 363:fnw255.

36. Watanabe K, Okuda M, Koga N. 2008. Newly developed system based on multiple enzyme restriction fragment length polymorphism-an application to proteolytic bacterial flora analysis, Soil Sci.Plant Nutr. 54:2, 204–215

37. Watanabe K, Koga N. 2009. Use of a Microchip Electrophoresis System for Estimation of Bacterial Phylogeny and Analysis of NO^3^ Reducing Bacterial Flora in Field Soils. Biosci. Biotech. Biochem. 73:479–488.

38. Watanabe K, Horinishi N, Matumoto K, 2015. Antibiotic-Resistant Bacterial Group in Field Soil Evaluated by a Newly Developed Method Based on Restriction Fragment Length Polymorphism Analysis. Adv. Microbiol. 5:807–816.

39. Watanabe K, Horinishi N, Matumoto K, Tanaka A, Yakushido K. 2015. Bacterial Groups Concerned with Maturing Process in Manure Production Analyzed by a Method Based on Restriction Fragment Length Polymorphism Analysis. Adv. Microbiol. 5:832–841.

40. Watanabe K, Horinishi N, Matumoto K, Sogabe Y. 2016. Identification and Enumeration Method of both Eukaryotic and Prokaryotic Microorganisms in Food Sample. Food Nutr. Sci. 7: 345–354.

41. Horinishi N, Matumoto K, Watanabe K. 2016. A Simple Evaluation System for Microbial Property in Soil and Manure. Adv. Microbiol. 6: 88–97.

42. Matumoto K, Shimada K, Horinishi, N, Watanabe K. 2016. Evaluation of the method based on restriction fragment length polymorphism analysis as simple analysis method of lactic acid bacteria in foods. Food Nutr. Sci. 7: 163–172.

43. Watanabe K, Horinishi N, Matsumoto K, Tanaka A, Yakushido K. 2016. A new evaluation method for antibiotic-resistant bacterial groups in environment. Adv. Microbiol. 6:133–151.

44. Weidner S, Arnold W, Puhler A. 1996. Diversity of Uncultured Microorganisms Associated with the Seagrass *Halophila stipulacea* Estimated by Restriction Fragment Length Polymorphism Analysis of PCR-Amplified 16S rRNAGenes. Appl. Environ.Microbiol. 62:766–771.

45. Cole JR, Chai B, Farris RJ, Wang Q, Kulam-Syed-Mohideen AS, McGarrell DM, Bandela AM, Cardenas E, Garrity GM, Tiedje JM. 2007. The Ribosomal Database Project (RDP-II): Introducing myRDP Space andQuality Controlled Public Data. Nucleic Acids Res. 35:D169–D172.

46. Nei M, Li WH. 1979. Mathematical Model for Studying Genetic Variation in Terms of Restriction Endonucleases. Proc. Natl. Acad. Sci. U.S.A. 76:269–5273.

47. Blodgett R. 2010. FDA, Bacterial Analytical Manual, Appendix 2: Most Probable Number from Serial Dilutions. http://www.fda.gov/Food/FoodScienceResearch/LaboratoryMethods/ucm109656.htm

48. Haas CN. 1989. Estimation of Microbial Densities from Dilution Count Experiments. Appl. Environ. Microbiol. 55:1934–1942.

49. Nubel U, Engelen B, Felske A, Snaidir J, Wieshuver A, Amann RI, Ludwig W, Backhaous H. 1996. Sequence Heterogeneities of Genes Encoding 16S rRNAs in *Paenibacillus polymyxa* Detected by Temperature Gradient Gel Electrophoresis J. Bacteriol. 178: 5636–5643.

50. Heuer H, Krsek M, Baker P, Smalla K, Wellington EH.. 1997. Analysis of Actinomycete Communities by Specific Amplification of Genes Encoding 16S rRNA and Gel-Electrophoretic Separation in Denaturing Gradients. Appl. Environ. Microbiol. 63:3233–3241.

51. Morimoto S, Hoshino Y. 2008. Methods for analysis of soil communities by PCR-DGGE (1) bacterial and fungal communities. Soil Microorg. 62:63–68.

